# Morphometry of newborn piglets and its relevance at weaning

**DOI:** 10.1101/2021.03.15.435415

**Authors:** Lucas M. Silva, Pedro H. S. Fidelis, Lígia V. L. Gomes, Gleyson A. Santos, Amanda M. A. Oliveira, Elias S. Medeiros, Marcelle S. Araújo, Rennan H. R. Moreira

## Abstract

This study aimed to evaluate the development of suckling piglets using morphometric parameters. Different models were created to predict the probability to occur any of the three weight classes (light, medium, and heavy) based on the piglets’ weaning weight. The variables in this research were birth weight (PW_B_), lactation length (Lac), and morphometric parameters– body length (BL), heart girth (HG), body mass index (BMI), ponderal index (PI), surface-mass ratio (SM), and birth order (BO). An adjustment of the ordinal regression was proposed to predict the weight classifications. The model with a significant effect of the Lac variables was selected. The light and heavy piglets, regardless of their morphometry, have a high chance of staying in the same weight class at weaning. However, this does not occur in medium piglets with diverse morphometry.

**Summary statement:** This manuscript demonstrates how morphometry can influence the development of animals, even with similar weights.

## Introduction

The genetic enhancement in hyperprolific sows has resulted in a significant increase in the number of piglets born by farrowing. As a consequence, piglets have lower birth weight and/or greater weight variability between them. The limitation of space within the sows’ uterus during gestation is one of the causes of this problem. This variability can have direct impacts on the mortality rate during the suckling period (Pinheiro and Dallanora, 2014). Usually, high mortality rates (11.5% to 18.6%) occur in the first seven days of life. The mortality of these piglets is one of the factors that reduce production on farms and jeopardize the financial performance (Aires et al., 2014).

Low birth weight is one of the main factors related to piglets’ mortality. However, deaths can also be related to their morphometry, since litter quality is associated not only to weight but also uniformity. In this case, morphological differences between animals in the same weight range can lead to diverse developments. Therefore, besides the piglet’s weight at birth, morphological characteristics (e.g., body mass index, ponderal index, surface-mass ratio) can be evaluation indicators for the positive performance of piglets in their successive life stages (Baxter et al., 2008, 2009). These indicators can be used to identify which animals will be below the average weight and potentially unprofitable early on (Huting et al., 2018).

The piglet’s morphometry is directly linked to its thermoregulation and can have a great impact on its viability. According to Herpin et al. (2002), lighter piglets have bigger body surface in relation to their weight and are, therefore, more prone to hypothermia. However, piglets with similar weight may have body surfaces in different sizes (Hales et al., 2013). Sometimes this distinction can indicate animals that are more likely to survive and/or develop better. Therefore, it can be affirmed that piglets with better body mass index and ponderal index also have better growth rate, ability to compete for mammary glands, and survival capacity.

In modern pig farming, it is essential to select in advance which piglets need special conditions in a given period, and mathematical models contribute to these predictions. With reliable and validated models, it is possible to estimate the pigs’ weight at slaughter and other zootechnical parameters (Silva et al., 2015). This study aimed to evaluate the development of suckling piglets based on their morphometry and determine the more accurate mathematical model to predict their weight class at weaning.

## Material and Methods

The procedures performed during the experiment followed the guidelines determined by the Committee on Animal Research and Ethics of the Universidade Federal Rural do Semiárido (registered under protocol no. 22/2020).

### Animals and Facilities

The experiment was performed on 30 hyperprolific (two to six farrowing) swine matrices (TN70) in lactation at the commercial farm located in the municipality of Croatá de São Gonçalo do Amarante, Ceará, Brazil.

The matrices were transferred from the gestation facility to different maternity facilities after 105 days of gestation. The gestation facilities had individual cages and a solid (concrete) floor. The maternity facilities are made up of partially slatted floors with a heated creep for the piglets.

### Evaluated parameters

One day after farrowing, the size of the litter was standardized at 12 piglets. The cutting and healing of the umbilical cord were performed shortly after birth. On the third day after farrowing, the teeth were clipped and the tail cut. Seven-day-old piglets were castrated.

Piglets were identified and weighed individually one day after farrowing. They were weighed again and had their morphometric measures (body length – BL; heart girth -HG) collected at weaning (20-days-old), as per Figure 1. The birth and weaning weights were measured using a 3 decimal place balance.

**Fig. 1.**
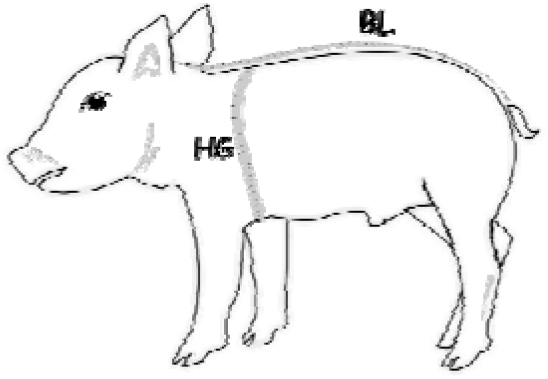
Piglet’s morphometric parameters: body length and heart girth (adapted from Dreamstime, 2020). (body length – BL; heart girth -HG).

The body length measurement started at the base of the ear going all the way until the first coccygeal vertebrae, following the midline suture of the cranium. Heart girth is the circumference measured right behind the forelimbs. Measurements were taken using a tape measure.

The body mass index (BMI) and ponderal index (PI) of all piglets were calculated based on their body length and birth weight (Amdi et al., 2013), as per the following equations:

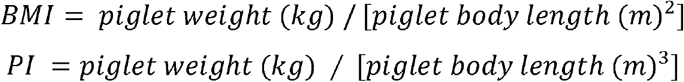

The correlation between the surface and mass (SM) was calculated using the formula proposed by Meeh (Brody et al., 1928):

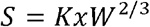

Where:

S: body surface area (dm^2^);

K: 0.07;

W: body weight (kg).

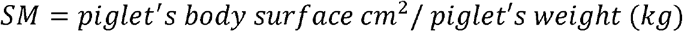

### Statistical analysis

In order to carry out an initial investigation of the data set, an exploratory analysis based on position measures (minimum, average, median, and maximum) and measures of variability (standard deviation and quantiles) was proposed.

Three classes were defined based on the normal distribution of the piglets’ weight at weaning (PW_W_): light (lighter than 3.967 kg), medium (3.967 to 5.095 kg), and heavy (heavier than 5.095 kg) piglets. Each class was determined with a 33.33% quartile. The average weight of the piglets was 4.531 kg (1.310 kg standard deviation), and the lactation length was 19.63 ± 1.41 days.

Ordinal regression was used to set a model capable of predicting (probability) which weight class is expected for the piglet at weaning. The weaning weight class was the dependent variable. The independent variables were the piglets’ birth weight (PW_B_), lactation length (Lac), and the morphometric parameters (BL, HG, BMI, PI, SM, and BO).

Based on the morphometric parameters and variables directly related to the piglet’s weight at weaning, the adjustment and comparison of the following models were suggested:

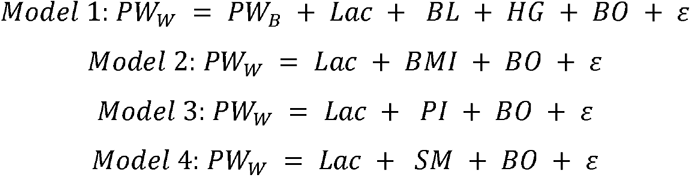

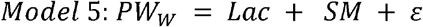

The models were verified by adjusting them according to the complete data set, which was divided into test data and training data.

1000 simple samples were extracted at random from 268 piglets for the training data, which represent 70% of the original data set (384 piglets). The remaining 116 piglets (test data) were left out of the analysis in order to verify the performance of the prediction (probability of each weight class to occur). Afterwards, the results were compared to the real weight class of the piglet at weaning (Figure 2).

**Fig. 2.**
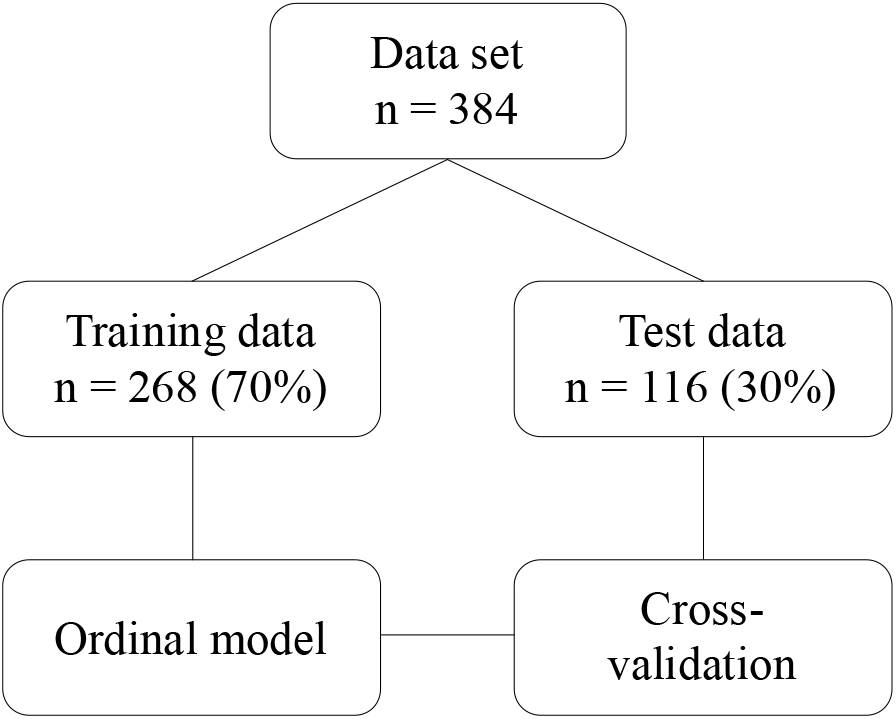
Flowchart of the adjustment in the ordinal regression model.

The most appropriate model was evaluated using a confusion matrix (Table 1), which establishes the correlation between the reference classes (observed weight classes) and the prediction (predicted weight classes for each model) by ordinal regression models. The “a” coefficient indicates the number of piglets that belonged to a reference class at weaning weight and that were correctly predicted to remain in the same class (reference = class and prediction = class). The “d” coefficient indicates the number of piglets that neither belong to the reference class nor were predicted to be in that class at weaning weight, which means a correct prediction by the model (reference = not class and prediction = not class). The “b” and “c” coefficients indicate the number of piglets incorrectly predicted by the models (reference = class and prediction = not class; reference = not class and prediction = class).

**Table 1.** Confusion matrix used to adjust the ordinal regression

| Prediction | Reference |  | Total |
| --- | --- | --- | --- |
|  | Class | Not class |  |
| Class | a | b | a+b |
| Not class | c | d | c+d |
| Total | a+c | b+d | a+b+c+d |

Class: number of weaning piglets belonging to one of the classes (light, medium, and heavy).
Not class: number of weaning piglets not belonging to one of the classes (light, medium, and heavy).

According to Jeune et al. (2018) and based on the confusion matrix, parameters to evaluate the accuracy of the models were obtained using sensitivity, precision, and Kappa’s values.

Sensitivity is the estimated probability (in percentage) of a correct prediction/result within the reference class (a/a+c) for each model. Precision is the likelihood that the model will provide correct results (a+d / a+b+c+d), which means that it is capable of predicting if the piglets will belong to a class when their reference is ‘class’ (same is true for ‘not class’). The value of Kappa can be classified as slight (0.00 to 0.20), reasonable (0.21 to 0.40), moderate (0.41 to 0.60), substantial (0.61 to 0.80), and almost perfect (0.81 to 1.00), according to Landis and Koch (1977).

After selecting the best models in cross-validation and based on the accuracy parameters, a single model was chosen using the AIC and BIC values. The best model was the one with the lowest AIC and BIC values

Based on the model that best described the piglets’ weight classes at weaning, the equations to obtain the probabilities of the piglet belonging to one of the three classes were presented.

All statistical analyses were performed using the R software (R Core Team, 2020), *ggplot2* (Wickham, 2016), and *MASS* (Venables and Ripley, 2002).

## Results

### Exploratory analysis

The evaluated parameters did not present a normal distribution (P<0.05), except for the piglet weight (P>0.05; Table 2).

**Table 2.** Descriptive statistics of the variables under study.

| Variables | Average | Standard Deviation | Minimum | PCTL (25%) | Median | PCTL (75%) | Maximum | p-SW |
| --- | --- | --- | --- | --- | --- | --- | --- | --- |
| PW<sub>B</sub> | 1.636 | 0.378 | 0.625 | 1.385 | 1.658 | 1.891 | 2.575 | 0.438 |
| PW<sub>W</sub> | 4.531 | 1.310 | 1.000 | 3.500 | 4.600 | 5.500 | 8.000 | 0.012 |
| BL | 26.79 | 2.40 | 19.00 | 25.00 | 27.00 | 29.00 | 32.00 | <0.001 |
| HG | 26.04 | 2.27 | 17.00 | 25.00 | 26.00 | 28.00 | 31.00 | <0.001 |
| BMI | 22.57 | 3.40 | 13.65 | 20.44 | 22.34 | 24.10 | 40.48 | <0.001 |
| PI | 84.82 | 14.74 | 47.77 | 76.02 | 82.68 | 91.66 | 192.74 | <0.001 |
| BSA | 965.89 | 152.16 | 511.70 | 869.76 | 980.39 | 1,070.53 | 1,315.07 | 0.044 |
| SM | 601.98 | 51.61 | 510.71 | 566.04 | 591.49 | 627.98 | 818.72 | <0.001 |
| Lac | 19.63 | 1.41 | 18.00 | 19.00 | 19.00 | 21.00 | 23.00 | <0.001 |
| BO | 2.70 | 1.49 | 1.00 | 1.00 | 2.00 | 3.20 | 6.00 | <0.001 |

PW<sub>B</sub>: piglet's birth weight; PW<sub>W</sub>: piglet's weaning weight; BL: body length; HG: heart girth; BMI: body mass index; PI: ponderal index; BSA: body surface area; SM: surface-mass ratio; Lac: lactation length; BO: birth order.

### Ordinal Regression

The median values for sensitivity were 66.67, 31.93, and 69.39% in Model 1; 68.29, 10.81, and 69.05% in Model 2; 64.86, 8.82, and 68.29% in Model 3; 65.88, 31.43, and 71.79 in Model 4; and 65.85, 32.35, and 70.83% in Model 5 for the weight classes 1, 2, and 3, respectively (Figure 3).

**Fig. 3.**
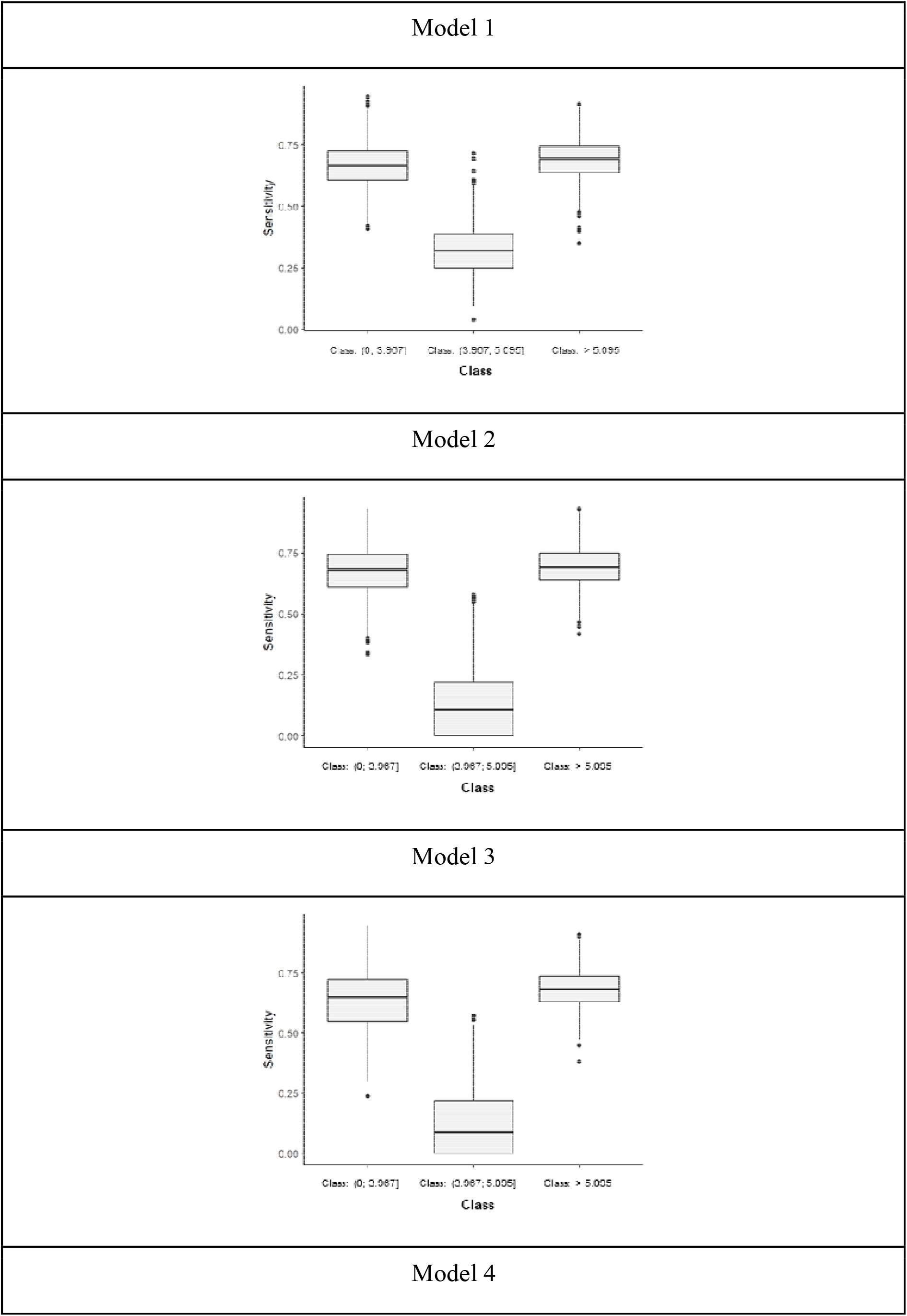
Evaluation of the performance of each model through cross-validation, represented by a box plot.

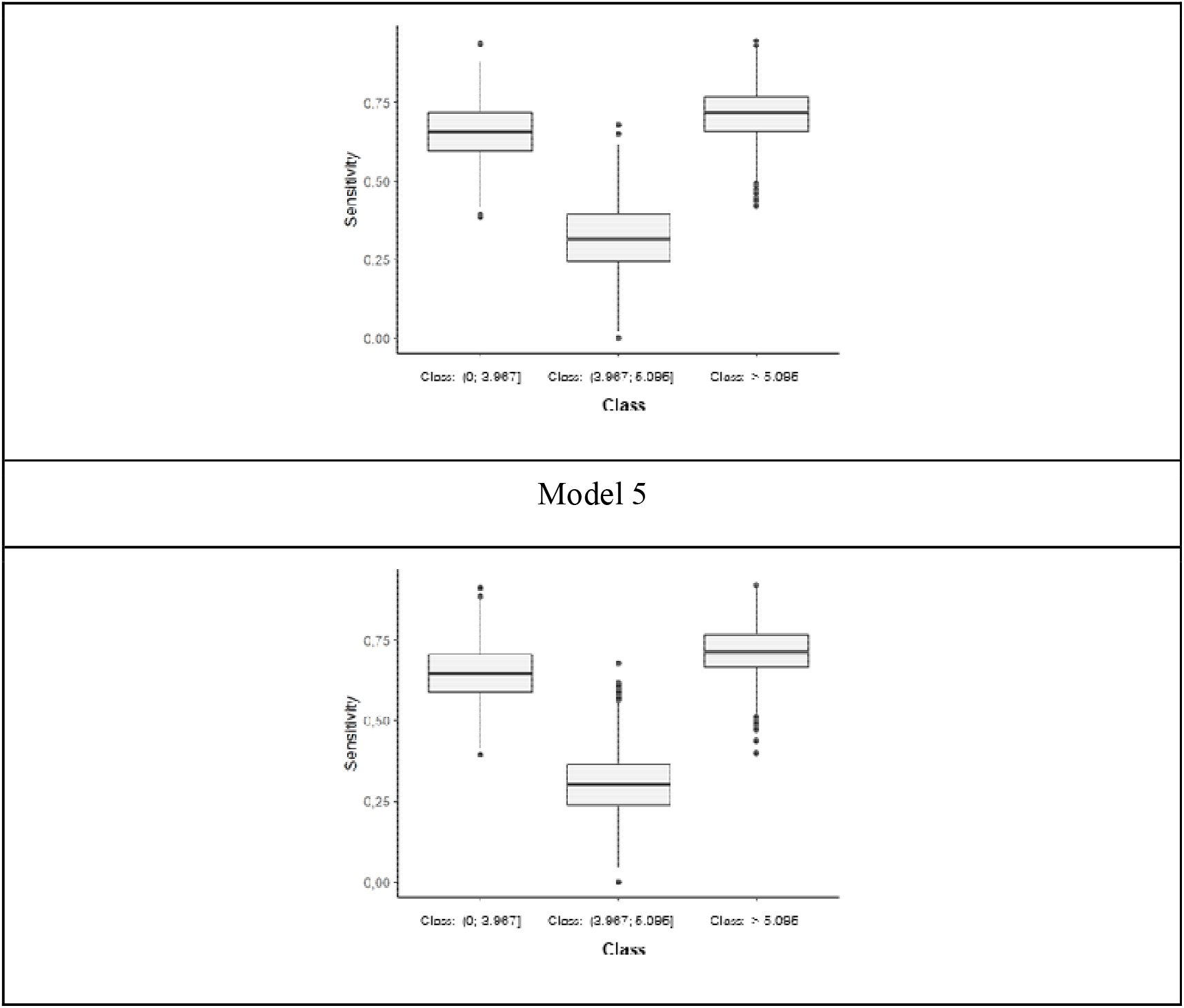

Models 2, 4, and 5 presented better sensitivities. However, the maximum difference observed between all models was only 2.08 percentage points. The models presented reasonable Cohen’s Kappa values, in which models 4 and 5 were higher than the other ones. In addition, both models had higher precision results in the evaluated set (Table 3).

**Table 3.**
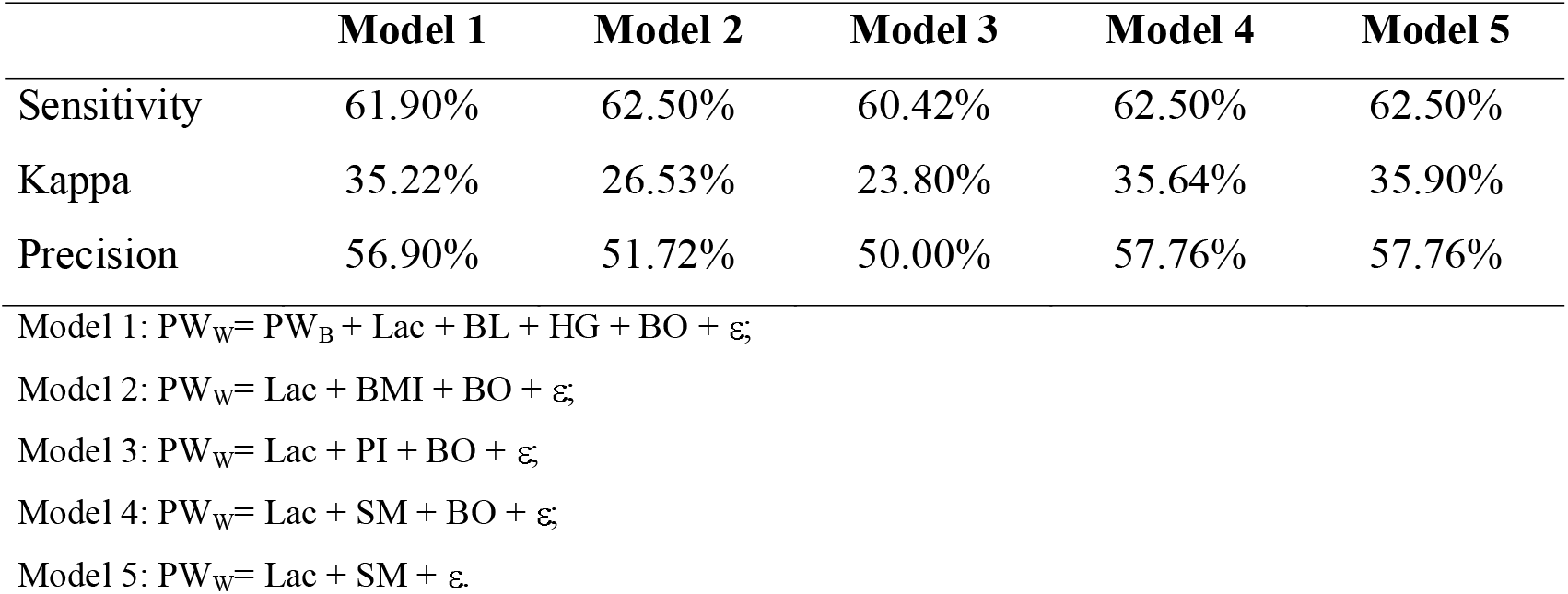
Median values obtained in the statistics analysis (sensitivity, Kappa, and precision) performed to evaluate the performance of each model through the cross-validation of the test set.

Models 4 and 5 presented greater sensitivity, Kappa, and precision. However, when analyzing the accuracy of these models, the covariate birth order was not statistically significant (P> 0.05) in Model 4. Thus, Model 5 was considered the best model to predict the probability of the piglets’ weaning weight. Comparing the models with better accuracy and considering the significance of the parameters for each model, the values of AIC and BIC reinforce Model 5 as the most appropriate since it presented the lowest values for these parameters (Table 4).

**Table 4.** Analysis of variance for the classification of weaning weight in Model 4.

| **Parameter** | **AIC** | **BIC** | **LR Chisq** | **Pr (>Chisq)** |
| --- | --- | --- | --- | --- |
| Model 4 | 690.226 | 709.979 |  |  |
| Lac |  |  | 52.172 | <0.001 |
| SM |  |  | 80.489 | <0.001 |
| BO |  |  | 0.517 | 0.472 |
| Model 5 | 688.743 | 704.546 |  |  |
| Lac |  |  | 55.675 | <0.001 |
| SM |  |  | 83.320 | <0.001 |

Lac: lactation length; SM: surface-mass ratio; BO: birth order.

Model 5 presented sensitivity results of 66.15% for the light class, 33.04% for medium, and 71.13% for heavy. It also had a Kappa value of 0.37 and precision of 58.30% (Table 5).

**Table 5.** Confusion matrix of the classification piglet’s weaning weight (PW_W_) developed using ordinal logistic regression – Model 5.

| **Expected** | **Reference** |  |  | **Total** |
| --- | --- | --- | --- | --- |
|  | (0 to 3.967] | (3.967 to 5.095] | > 5.095 |  |
| (0 to 3.967] | 86 | 37 | 13 | 136 |
| (3.967 to 5.095] | 23 | 37 | 28 | 88 |
| > 5.095 | 21 | 38 | 101 | 160 |
| **Total** | 130 | 112 | 142 | 384 |

The equations used to calculate the probabilities of belonging to any of the three classes are shown in Table 6.

**Table 6.** Probability estimates for each weight class in Model 5 Class Probability

| Class | Probability |
| --- | --- |
| Light | $P_1 = \frac{1}{1 + e^{-2.018 - (0.593 \times Lac - 0.021 \times SM)}}$ |
| Medium | $P_2 = \frac{1}{1 + e^{-0.346 - (0.593 \times Lac - 0.021 \times SM)}} - P_1$ |
| Heavy | $P_3 = 1 - P_2 - P_1$ |

Based on Model 5 and analyzing the parameters one at a time, it can be observed that as the value of the surface-mass ratio increases, the probability of the piglet belonging to the light class at weaning also increases. A different result is observed at the lactation length, in which older piglets have a higher probability of being in heavier classes at weaning (Figure 4).

**Figure 4.**
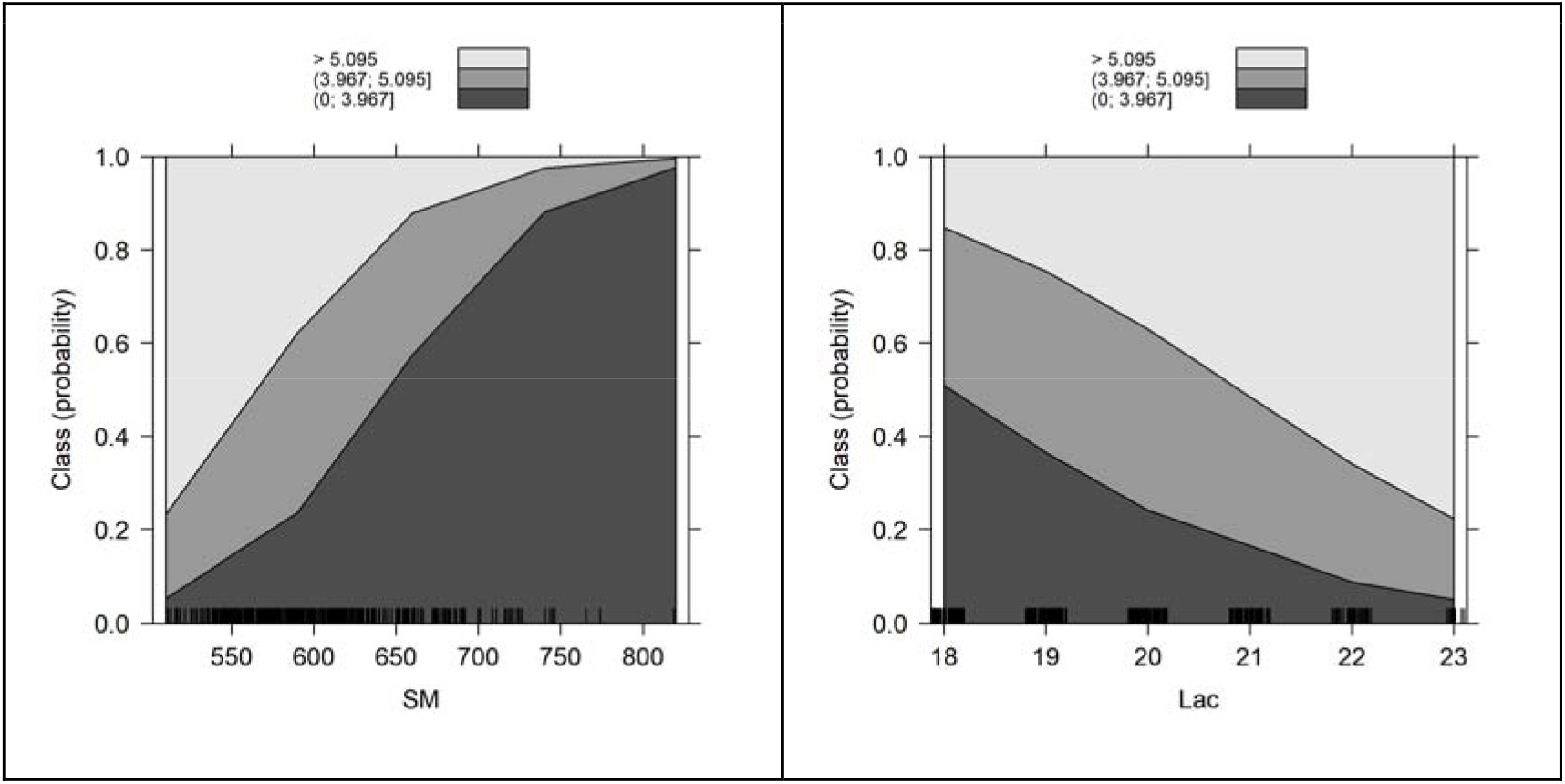
Probabilities of the piglets belonging to any weight class in each parameter (surface-mass ratio – SM; and lactation length – Lac) of Model 5.

## Discussion

### Ordinal regression models

The ordinal regression models allowed the identification of important variables at birth and also to estimate the weight classes of the piglets at weaning. Greater accuracy was observed in Model 5, which had significant results for the variables lactation length and surface-mass ratio. It is worth mentioning that these variables implicitly represent all other variables (BMI and PI) since surface-mass ratio (SM) is a response variable originated from implicit analytic parameters.

Birth order is directly related to the weight at weaning, and it influences the number of piglets born and their birth weight. However, this variable did not have a significant influence on the prediction of the tested models (Pinheiro et al., 1996) and invalidated the models 1, 2, 3, and 4.

All models had low accuracy in predicting the weight of piglets at weaning in the medium weight class, which means that they have low sensitivity rates. Based on this information, it can be inferred that the morphometry of piglets with medium birth weight has a low influence on their weight at weaning. Therefore, these piglets may present different development. This is an important discovery, as it demonstrates the need to improve the conditions for fetal development, especially the sows’ nutrition. Through nutrition, it is possible to modulate the development and improve the morphometry of piglets at birth. Thus, piglets born with medium weight remain in this class at weaning, or may even end up reaching the heavy class.

On the other hand, Model 5 showed a high sensitivity for the weight classes light and heavy, indicating that the morphometry has a strong influence on the weight of the piglets at weaning. If the piglets are born light or heavy, regardless of their morphometry, they are more likely to remain in their respective weight classes at weaning. Greater management, nutrition, and environmental conditions are necessary for light piglets, in order to reduce that their mortality rate during lactation and also to avoid the transmission of pathogens.

Model 5 was chosen because it has good sensitivity, reasonable Kappa, 58.3% precision, and lower AIC and BIC values. In addition, this model is easier to apply in farms, since lactation length and surface-mass ratio are parameters that are easy to measure. According to Pozza et al. (2008), the prediction models must have simple measurement parameters so they could be used in the field.

### Morphometric Parameters

Birth weight is one of the main factors related to piglet survival. The lack of uniformity contributes to a higher occurrence of light piglets in the litter. Piglets that are light at birth have fewer energy reserves in their body and need more time to feed on their mother’s milk (Panzardi et al., 2009). Therefore, they take longer to gain weight and, consequently, need more time to be weaned.

In addition to birth weight, the morphometry influences the development of piglets at weaning, in which those with a high surface-mass ratio are the ones most likely to belong to class 1. Light animals with a body that has a large surface area are more prone to suffer from cold temperatures (due to heat losses). Also, they are less competitive and more susceptible to mortality. Energy reserves and thermoregulation are relevant aspects in the early stages of life, and their deficiencies may compromise the piglet’s development (Alonso-Spilsbury et al., 2007).

The early identification of animals with deformed morphometry is seen as an important strategy. This allows professionals to plan the most effective way to alleviate the problems arising from piglets with delayed intrauterine growth. The morphometric parameters used in the models are considered to be excellent predictors of the animal’s development. In addition to indicating the piglets’ ability to survive from their birth until weaning, morphometry can also be an important strategy for future development assessments (Litten et al., 2005).

## Conclusion

Light and heavy piglets, regardless of their morphometry, have a high chance of staying in the same weight class at weaning. However, this does not occur in medium piglets with different morphometry.

Further studies should be carried out in order to improve the morphometry of light piglets, increasing their chances of survival and future development.

In addition to weight, the results indicate that morphometric parameters are fundamental to evaluate the development of piglets.

## Acknowledgment

The authors want to acknowledge the Regina Farm and Marcus Vinícius Cardoso Trento.

